# A polyploid admixed origin of beer yeasts derived from European and Asian wine populations

**DOI:** 10.1101/466276

**Authors:** Justin C. Fay, Ping Liu, Giang T. Ong, Maitreya J. Dunham, Gareth A. Cromie, Eric W. Jeffery, Catherine L. Ludlow, Aimée M. Dudley

## Abstract

Strains of *Saccharomyces cerevisiae* used to make beer, bread and wine are genetically and phenotypically distinct from wild populations associated with trees. The origins of these domesticated populations are not always clear; human-associated migration and admixture with wild populations have had a strong impact on *S. cerevisiae* population structure. We examined the population genetic history of beer strains and find that ale strains and the *S. cerevisiae* portion of allotetraploid lager strains were derived from admixture between populations closely related to European grape wine strains and Asian rice wine strains. Similar to both lager and baking strains, ale strains are polyploid, providing them with a passive means of remaining isolated from other populations and providing us with a living relic of their ancestral hybridization. To reconstruct their polyploid origin we phased the genomes of two ale strains and found ale haplotypes to both be recombinants between European and Asian alleles and to also contain novel alleles derived from extinct or as yet uncharacterized populations. We conclude that modern beer strains are the product of a historical melting pot of fermentation technology.

## Introduction

The brewer’s yeast *Saccharomyces cerevisiae* is known for its strong fermentative characteristics. The preference for fermentation in the presence of oxygen arose as a multi-step evolutionary process around the time of an ancient genome duplication, endowing numerous species with the ability to produce levels of ethanol toxic to many microorganisms (Dashko et al., 2014; Hagman et al., 2013). One of these species, *S. cerevisiae*, also gained the ability to competitively dominate many other species in high sugar, low nutrient environments, such as grape must (Williams et al., 2015). Wine is largely fermented by *S. cerevisiae* and is thought to be the first fermented beverage, having been made for over 9,000 years (McGovern, 2013). However, *S. cerevisiae* is not the only *Saccharomyces* species used to make fermented beverages; others, particularly *S. uvarum*, *S. kudriavzevii*, *S. eubayanus* and hybrid derivatives are also used, particularly for fermentations at low temperatures (Almeida et al., 2014; González et al., 2006; Libkind et al., 2011; Sipiczki, 2008). Besides *S. cerevisiae*, the most widely used species is *S. pastorianus*, an allopolyploid hybrid of *S. cerevisiae* and *S. eubayanus*, used to make lager beer (Libkind et al., 2011). The use of this hybrid emerged during the 15th century in Europe and was formed from a *S. eubayanus* strain closely related to wild populations from North America and Tibet (Bing et al., 2014; Peris et al., 2016) and a *S. cerevisiae* strain, related to those used to ferment ales (Dunn and Sherlock, 2008; Gallone et al., 2016; Gonçalves et al., 2016). The origin of ale and other domesticated strains of *S. cerevisiae* is beginning to emerge through comparison with wild populations (Duan et al., 2018; Gallone et al., 2016; Gonçalves et al., 2016; Legras et al., 2014; Peter et al., 2018).

Multiple genetically distinct populations of *S. cerevisiae* have been found associated with fermented foods and beverage. These include grape wine, Champagne, sake and rice wine, palm wine, coffee, cacao, cheese, and leavened bread (Duan et al., 2018; Fay and Benavides, 2005; Legras et al., 2007, 2018; Ludlow et al., 2016). Ale strains have also been found to be both genetically and phenotypically differentiated from other strains (Gallone et al., 2016; Gonçalves et al., 2016). However, the origin of such domesticated groups is not always clear as it requires comparison to wild populations from which they were derived and these wild populations have not all been identified. The best characterized wild populations of *S. cerevisiae* have been isolated from oak and other trees in North America, Japan, China and Europe (Almeida et al., 2015; Hyma and Fay, 2013; Sniegowski et al., 2002; Wang et al., 2012), the latter of which is most closely related to and the presumed wild lineage from which European wine strains were derived.

Despite clear differences among many domesticated groups, human-associated admixture is common (Cromie et al., 2013; Hyma and Fay, 2013; Ludlow et al., 2016; Tilakaratna and Bensasson, 2017) and can blur the provenance of domesticated strains. For example, wine strains show a clear signature of admixture with other populations, and clinical strains appear to be primarily derived from admixed wine populations (Liti et al., 2009; Schacherer et al., 2009; Strope et al., 2015). Ale strains, with the exception of a few found related to sake and European wine lineages, have no obvious wild population from which they were derived (Gallone et al., 2016; Gonçalves et al., 2016).

In this study we examined the origin of ale and lager strains in relation to a diverse collection of *S. cerevisiae* strains. Through analysis of publicly available genomes and 107 newly sequenced genomes, we inferred a hybrid, polyploid origin of beer strains derived from admixture between close relatives of European and Asian wine strains. This admixture suggests that early industrial strains spread with brewing technology to give rise to modern beer strains, similar to the spread of domesticated plant species with agriculture.

## Results

We sequenced the genomes of 47 brewing and baking strains and 65 other strains of diverse origin for reference. Combining these with 430 publicly available genomes we found nearly all the brewing strains closely related to previously sequenced ale and lager strains (Figure S1). Through analysis of population structure we identified 13 populations, four of which contain the majority (64/76) of beer strains. The four beer-associated populations consisted of predominantly lager strains, German ale strains (Ale 1), British ale strains (Ale 2) and a mixture of beer and baking strains (Beer/baking), and are consistent with previously identified groups of beer strains (Gallone et al., 2016; Gonçalves et al., 2016). The remaining populations were similar to previously characterized groups (Almeida et al., 2015; Liti et al., 2009; Ludlow et al., 2016) and were classified by the most common source and/or geographic region of isolation as Laboratory, Clinical, Asia/sake, Europe/wine, Mediterranean/oak, Africa/Philippines, China/Malaysia, and two populations from Japan/North America (Table S2).

To identify the most likely founders of the four beer populations we used a composite likelihood approach to infer population relationships while accounting for admixture, which can obfuscate population phylogenies (Pickrell and Pritchard, 2012). The inferred admixture graph grouped the four beer populations together, with the lager and two ale populations being derived from the lineage leading to the Beer/baking population (Figure 1). The four beer populations are most closely related to the Europe/wine population. However, the admixture graph also showed strong support for two episodes of gene flow into the beer lineages resulting in 40-42% admixture with the Asia/sake population. We confirmed these admixture events using *f_4_* tests for discordant population trees, which are caused by admixture (Peter, 2016; Reich et al., 2009). All *f_4_*(Europe, test; Asia, Africa) statistics were significant for tests of the four beer populations (Z-scores < −21.3), whereas the *f_4_*(Europe, Mediterranean; Asia, Africa) statistic was much closer to zero for the Mediterranean/oak population (Z-score = 3.4). Similar results were obtained using the China or Japan/North American populations rather than Africa (Table S3). Thus, the beer populations were derived from admixture events between a population closely related to the Europe/wine population and Asia/sake population. Consistent with prior studies (Ludlow et al., 2016; Strope et al., 2015), we also found admixture events between the Europe/wine population and both the lab and clinical population.

**Figure 1.**
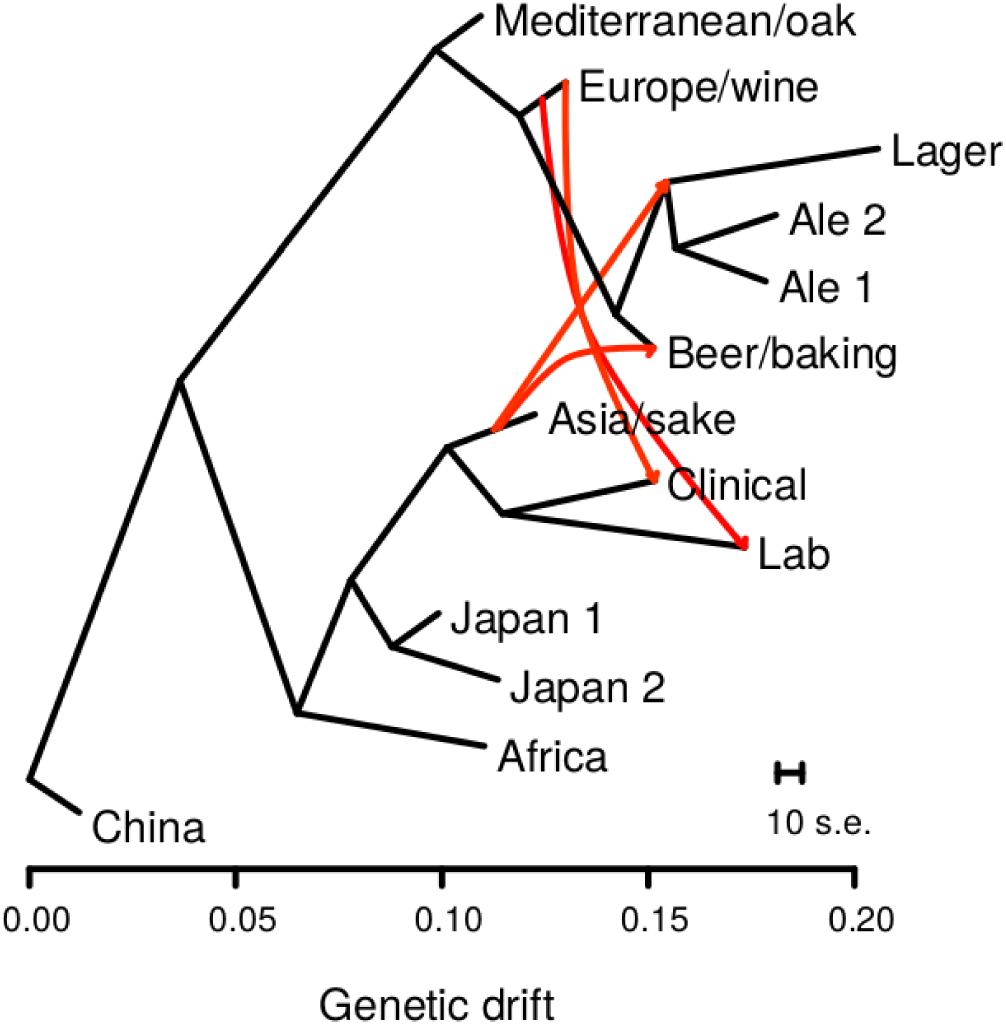
Admixture graph of population relationships shows admixture events from the Asia/sake to multiple beer populations. Population relationships were inferred using TreeMix and horizontal branch lengths are proportional to genetic drift with the scale bar showing the average of ten standard errors of the sample covariance matrix. Red arrows show admixture events with migration weights over 0.40, indicating the fraction of alleles derived from a source population. The migration from the ancestor of the Mediterranean/oak to the Clinical population is not shown for clarity.

To quantify the degree of admixture for each of the beer strains we calculated *f_4_* admixture proportions using the Europe/wine and Asia/sake populations (Green et al., 2010; Peter, 2016). For the 64 beer strains we estimated 36.7-46.7% of their genome was derived from the Asia/sake population, with an average of 39.6%. The high proportion yet narrow range of admixture implies little to no subsequent back-crossing following admixture.

Polyploidy is a known mechanism of reproductive isolation (Mallet, 2007), and could facilitate isolation of the beer populations. In yeast, tetraploid x diploid crosses produce triploids, and triploids suffer from high rates of spore inviability due to aneuploidy (Campbell et al., 1981; Loidl, 1995). Polyploidy is enriched in beer and baking strains (Gallone et al., 2016; Legras et al., 2007; Peter et al., 2018), and the Beer/baking population includes 10 previously studied strains, all of which were found to be triploid or tetraploid and to exhibit high rates of heterozygosity (Ludlow et al., 2016). These strains were previously found to group with other strains isolated from diverse sources around the world (Pan/Mixed 2 in Ludlow et al. 2016). To identify triploids and tetraploids strains, we used the expected allele frequency at heterozygous sites: 50% for diploids, 33 and 66% for triploids, and 25, 50, and 75% for tetraploids (Figure 2). We note that this approach does not resolve ploidy unless there are a sufficient number of heterozygous sites with high enough read depth to estimate allele frequencies. Nevertheless, out of 105 strains with an abundance of heterozygous sites, we identified 15 diploid, 23 triploid and 28 tetraploid strains (Figure S2). The remaining 39 strains did not have sufficient read depth or exhibited dispersed allele frequencies with no clear indication of ploidy level. Of the 51 polyploid strains (N>2), 45 (88%) were in one of the four beer populations, of which 29 were beer strains and six were baking strains. The remaining ten polyploids assigned to the beer populations include three isolates from green coffee beans and were previously assigned to a Pan/Mixed 2 population (Ludlow et al., 2016), a group of predominantly human-associated strains.

**Figure 2.**
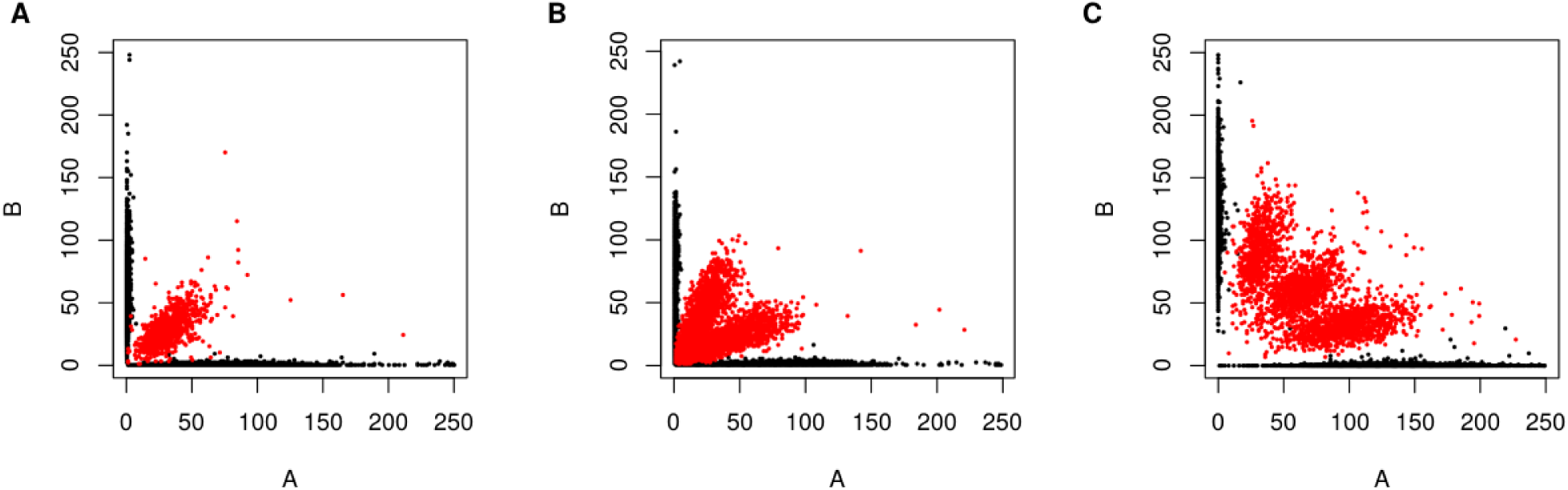
Diploid, triploid and tetraploids are distinguished by read counts at heterozygous sites. Plots show examples of a diploid (A, YO700), triploid (B, TUM205) and tetraploid (C, YMD1952) strain by the number of reads with the reference allele (A) versus the alternative allele (B) for heterozygous (red) and homozygous (black) sites.

Given the admixed origin of beer strains, we wanted to know which populations the heterozygous sites indicative of polyploidy were derived from. We examined heterozygous sites in the beer populations in relation to other strains by clustering SNPs and grouping strains by their inferred population membership (Figure 3). Excluding the lager population, where are not heterozygous, the three beer populations are predominantly heterozygous for alleles abundant within either the Asia/sake or Europe/wine population. However, all four beer populations also carry alleles not present in any of the other populations. The presence of heterozygous, beer-specific alleles suggests that these alleles were derived from admixture between a lineage closely related to the Europe/wine lineage and/or the Asia/sake lineage. The large number of beer-specific alleles is unlikely to have accumulated in the recent past since the formation of the polyploids. The beer groups have between 6,558 and 13,728 alleles present at 25% frequency or more in the group but not in any other population. Using these beer-specific alleles we found divergence at four-fold degenerate synonymous sites was 0.153, 0.100, 0.087, and 0.069% along the Ale 1, Ale 2, Beer/baking and Lager lineages, higher than expected to have accumulated since the use of these strains for brewing purposes (see Discussion) and not much less than the rate of divergence between the Europe/wine and Asia/sake population (0.592%). Combined with the observation that these variants are mostly heterozygous, we infer that they were present when the strains became polyploid and originated from an extinct or as yet to be characterized yeast population related to the Europe/wine and/or Asia/sake population.

**Figure 3.**
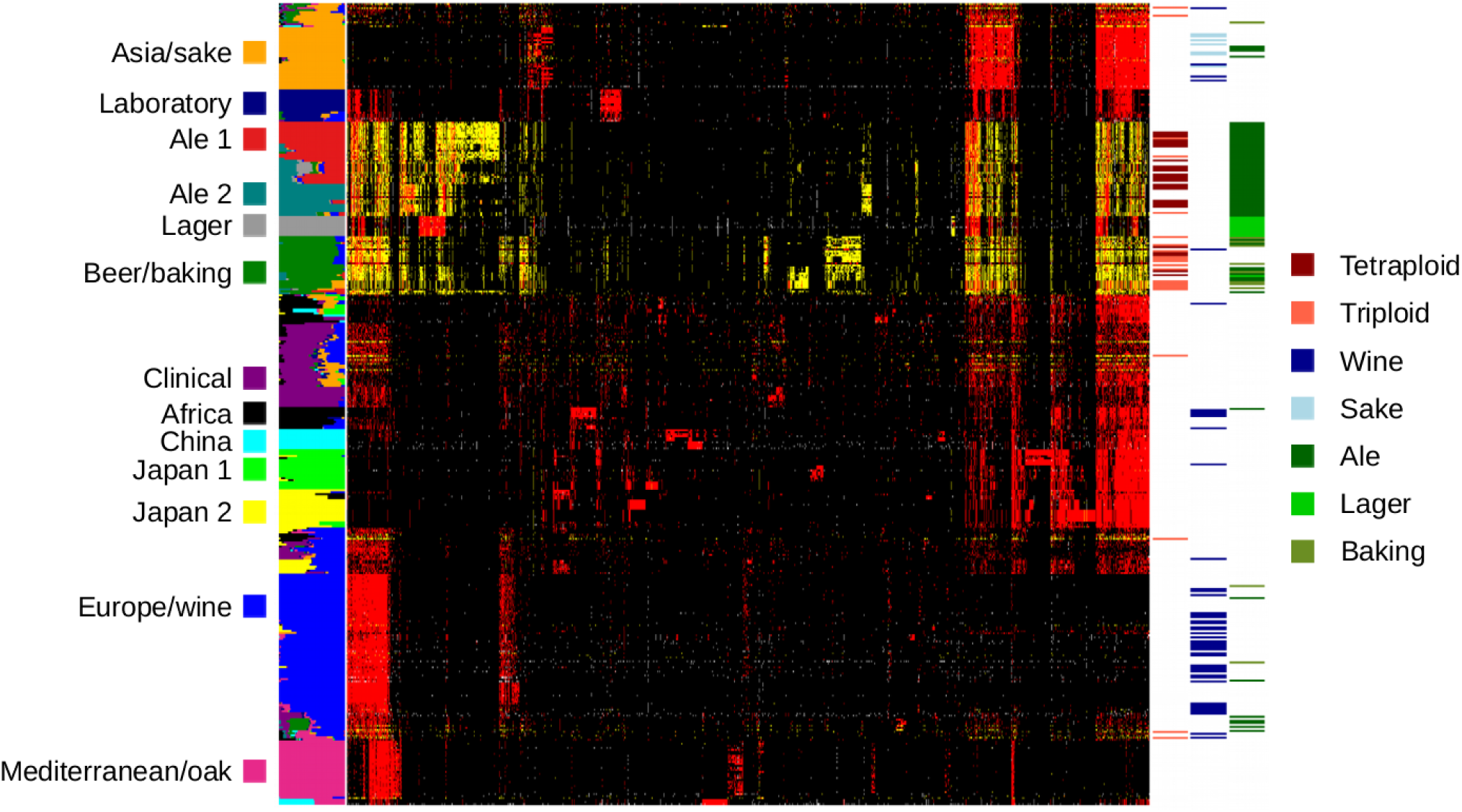
Beer populations are heterozygous for alleles shared with Europe/wine and Asia/sake populations. Genotypes of 2000 randomly selected SNPs are shown for 339 strains grouped by their ancestry to 13 populations shown by the colored panel on the left. Genotypes are homozygous for the major allele (black), minor allele (red), or heterozygous (yellow) and SNPs (columns) were ordered by hierarchical clustering. The panel on the right indicates triploid and tetraploid strains, grape and sake wine strains, and ale, lager and baking strains.

The presence of heterozygous European, Asian and beer-specific alleles enabled us to use haplotype phasing to test whether there has been recombination between European and Asian haplotypes and whether beer-specific alleles reside on European or Asian haplotypes. We used long-read sequencing to phase two beer strains: a German ale strain in the Ale 2 population (A.2565) and a Belgian ale strain in the Beer/baking population (T.58). Both strains were inferred to have over 99% ancestry to their assigned population, and both are likely polyploids (Table S2). As a control we phased the genome of a hybrid (YJF1460) generated between a Europe/wine strain and a Japan/North America 2 oak strain. Because of uncertainty in the ploidy as well as the possibility of variable ploidy levels (aneuploidy) across the genome, we developed a phasing algorithm that merges reads into consistent haplotypes and makes no assumptions about ploidy.

Phasing of the three strains yielded predominantly two haplotypes in the YJF1460 control and three or four haplotypes in the two ale strains (Figure 4 and Figure S4). The majority of phased haplotypes in the two ale strains carried a mixture of European and Asian alleles. In contrast, the YJF1460 control showed few haplotypes with both European and Asian alleles, indicative of haplotype switching errors or mitotic exchange. Eliminating regions of haplotype switching less than 4 kb in length, which could result from genotype errors or mitotic gene conversion, we counted the number of switches within the phased haplotypes between European and Asian alleles and found 12 in YJF1460, 346 in T.58 and 199 in A.2565 (Table S4). Consequently, most haplotypes present in the two ale strains represent recombinant haplotypes as opposed to pure European-related or Asian-related haplotypes (Figure 4 and Figure S4); only 22% of the T.58 genome and 19% of the A.2565 genome carried haplotypes with over 95% European or 95% Asian alleles. In contrast, 88% of the YJF1460 genome was inferred to carry pure European or Asian haplotypes. While it is difficult to know how much recombination occurred prior to polyploidy and how much occurred subsequently through mitotic recombination or gene conversion, mitotic events have contributed to diversification of beer strains subsequent to polyploidy; there are four large and a number of smaller regions in A.2565 that exhibit loss of heterozygosity (Figure 4). While the fixed haplotypes in these regions are also recombinants of European and Asian derived haplotypes, loss of heterozygosity is a distinct signature of mitotic recombination.

**Figure 4.**
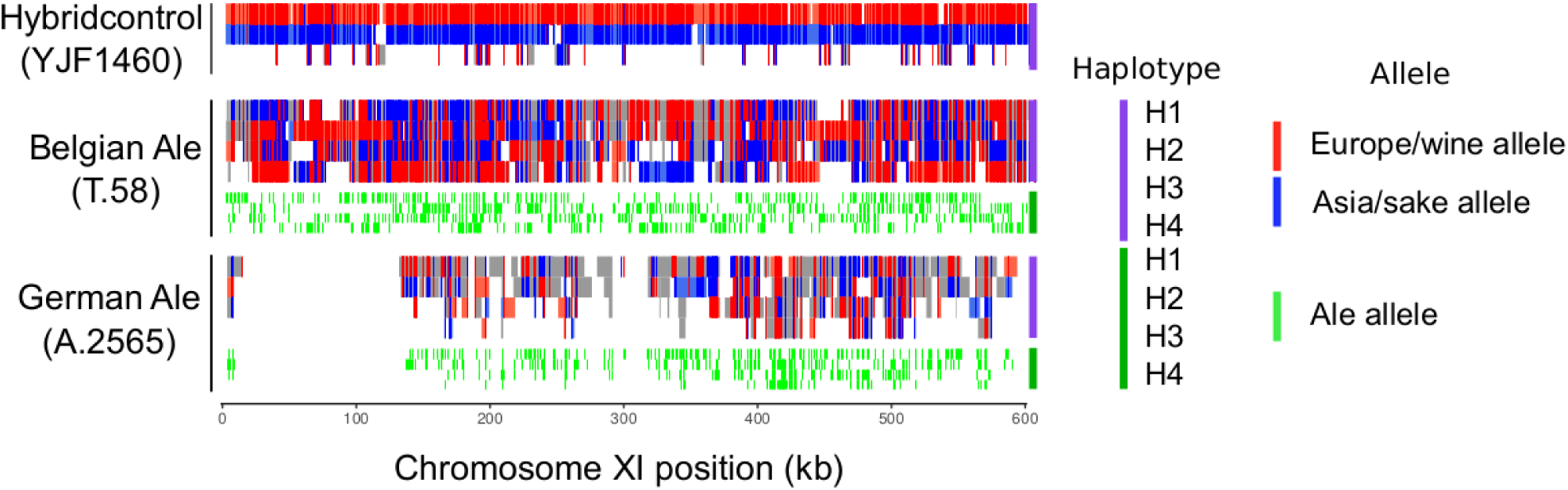
Phased haplotypes show recombination between European and Asian alleles. Panels for two ale strains (T.58 and A.2565) and the control hybrid (YJF1460) show haplotypes and allele configurations across chromosome XI. The purple panel shows haplotypes with more than 95% European or Asian alleles are shown in red and blue, respectively, and in gray otherwise. Ticks indicate European (red) and Asian (blue) alleles. Haplotypes were assigned labels H1-H4 in order of longest to shortest, except for YJF1460 where they were assigned based on predominance of Europe/wine (H1, red) or Asia/sake (H2, blue) alleles. The green panel shows Ale-specific alleles for the four haplotypes (H1-H4) by green ticks. The first 100 kb of A.2565 shows a region exhibiting loss of heterozygosity.

The polyploid beer strains also carry beer-specific alleles not present in other strains. These beer-specific alleles could have been inherited from an ancestral population that split from the European/wine, Asian/sake population or both. To distinguish between these possibilities we examined the distribution of beer-specific alleles on the phased haplotypes and found most (~80%) were on mixed haplotypes, having at least 5% European and 5% Asian alleles. The remaining beer-specific alleles were equally distributed between predominantly European or predominantly Asian haplotypes (Table S4). Because many of the beer-specific alleles are not shared between the Ale 1, Ale 2, Beer/baking and Lager populations, we can infer multiple origins of the four beer populations despite similar episodes of admixture and polyploidy, demonstrating that the *S. cerevisiae* contribution to lager strains did not simply come from one of the other beer strain lineages.

## Discussion

Inferring the origin of domesticated organisms can by complicated by extinction of wild progenitor populations, human-associated migration, polyploidy and admixture with wild populations. In this study we find that extant beer strains are polyploid and have an admixed origin between close relatives of European and Asian wine strains. Ale genomes, like lager genomes, carry relics of their parental genomes captured in a polyploid state as well as novel beer alleles from an extinct or undiscovered population. Loss of heterozygosity through mitotic exchange provided a means of strain diversification but has also potentially eroded precise inference of the timing and order of events giving rise modern beer strains. Below we discuss models and implications for an admixed, polyploid origin of beer strains.

Polyploidy is thought to mediate rapid evolution (Van de Peer et al., 2017) and is common in domesticated plants (Salman-Minkov et al., 2016). Prior work showed that polyploidy is common in beer and baking strains (Gallone et al., 2016; Legras et al., 2007; Peter et al., 2018). We find that the Ale 1, Ale 2 and Beer/baking population all have a polyploid origin. Although not all strains had sufficient coverage for calling polyploidy, all those that did were either triploid or tetraploid. Chromosome level aneuploidy is also more common in strains within the Ale 1 (52%), Ale 2 (19%) and Beer/baking (52%) populations than in the non-beer populations (5.1%). A notable consequence of both polyploidy and aneuploidy is that they can limit admixture with haploid or diploid strains due to low spore viability (Campbell et al., 1981; Loidl, 1995), thereby maintaining their brewing characteristics. Indeed, beer strains exhibit low sporulation efficiency and spore viability (Gallone et al., 2016). Interestingly, both grape wine and particularly sake wine strains have also evolved more limited capacities to interbreed through low sporulation efficiencies (Gerke et al., 2009; Nakazawa et al., 1992).

Human-associated admixture is well documented in wine strains, which have been dispersed around the globe with the spread of viticulture (Cromie et al., 2013; Hyma and Fay, 2013; Ludlow et al., 2016; Tilakaratna and Bensasson, 2017). However, admixture between close relatives of European grape wine and Asian rice wine populations presents a conundrum regarding where and how these populations became admixed. A crucial yet unresolved piece of information is where European wine strains were domesticated. The discovery of a Mediterranean oak population closely related to European wine strains suggests a European origin of wine strains (Almeida et al., 2015). However, copper and sulfite resistance, common in European wine strains, is also common in strains isolated from oak trees outside of vineyards in Slovenia (Dashko et al., 2016). An alternative model is that the Mediterranean oak population is either a feral wine population or both the European wine and Mediterranean oak populations are non-native. Analysis of a diverse collection of Asian strains suggested an East Asian origin of all domesticated *S. cerevisiae* strains, including European wine strains (Duan et al., 2018). Domestic populations from solid and liquid state fermentations (bread, milk, distilled liquors, rice and barley wines) were found related to wild populations from East Asia. In support of European wine and Mediterranean oak populations also originating in East Asia, these populations carried duplicated genes involved in maltose metabolism and grouped with fermented milk and other strains isolated from China. However, this model also has some uncertainty given the small number of Chinese isolates within the European wine group, the dispersion of European wine strains with viticulture, and the absence of samples from the Caucasus where grapes are thought to have been domesticated (McGovern, 2013; Myles et al., 2011).

Considering the uncertainty in where European wine strains were domesticated, we put forth two hypotheses regarding the admixed origin of beer strains. First, European wine strains were domesticated in East Asia and admixed *in situ* with a population related to the Asia/sake group that we describe, which contains eight sake/rice wine strains, seven distillery strains and seven bioethanol strains, mostly from Asia. Second, European wine strains were domesticated in Europe from a Mediterranean oak population, or perhaps in the Caucasus, and the admixed beer populations arose through East-West transfer of fermentation technology including yeast by way of the Silk Route. Resolving these scenarios would be greatly facilitated by finding putative parental populations of diploid but not necessarily wild strains that carry alleles we find to be unique to the Ale 1, Ale 2, Beer/baking and Lager groups. As yet, such populations have not been sampled or are extinct.

Even with a clear signature of a polyploid and admixed origin of beer strains, there are uncertainties regarding the founding strains and the order of events. In principle, if each beer genome remained static after polyploidy, then the number of polyploidy events (founders) and the number of generations of admixture prior to polyploidy could be determined. However, polyploid genomes are often labile and it is hard to know the extent to which mitotic recombination and gene conversion have altered genetic variation in the beer populations. In yeast, the rate of mitotic gene conversion and recombination has been estimated to be 1.3 × 10^-6^ per cell division and 7 × 10^-6^ per 120 kb, respectively (Lee et al., 2009; Yim et al., 2014), and both can lead to loss of heterozygosity. Converting to the size of a tetraploid genome (~48 Mbp) we expect 0.0038 (using a median track length of 16.6 kb) conversion events and 0.0028 recombination events across the genome per cell division. Three lines of evidence support the role of these mitotic events in beer strains. First, many of the switches between the European and Asian alleles involved one or a small number of adjacent SNPs rather than long segments, indicative of gene conversion (Table S4). Second, one strain (A.2565) shows clear loss of heterozygosity on multiple chromosomes, indicative of mitotic recombination (Figure S4). Third, there is substantial genotype diversity within each of the beer populations (Figure 3). This would be expected to occur if loss of heterozygosity occurred during strain divergence but subsequent to the founding of each beer population.

Two other factors besides mitotic gene conversion and recombination must be considered in regards to diversity within the beer populations: outcrossing and *de novo* mutation. Outcrossing with strains outside of the beer population is unlikely since there is no evidence for this type of admixture in our analysis and admixture proportions from the Asian population is fairly constant at 37-47% across beer strains. However, it is worth noting that outcrossing of strains within or between different beer populations may not easily be detected. *De novo* mutations have undoubtedly occurred, but even using a reasonable estimate of 150 generations per year for brewing strains (Gallone et al., 2016) and a per base mutation rate of 5×10^-10^ (Lang and Murray, 2008), the beer lineage substitution rates yield divergence times of 2.0×10^4^ (Ale 1), 1.3×10^4^ (Ale 2), 1.1×10^4^ (Beer/baking) and 9.2×10^3^ (Lager) years. Thus, a sizable fraction of beer-specific alleles was likely inherited from populations closely related to European wine and Asian wine populations rather than *de novo* mutations that accumulated subsequent to polyploidy. Regardless of the relative impact of mitotic recombination, gene conversion, outcrossing and *de novo* mutation, beer strains have diversified from one another but have remained relatively distinct from other populations of *S. cerevisiae* (Gallone et al., 2016; Gonçalves et al., 2016).

In conclusion, beer strains are the polyploid descendants of strains related to but not identical to European grape wine and Asian rice wine strains. Thus, similar to the multiple origins of domesticated plants, including barley (Poets et al., 2015) and rice (Choi et al., 2017; Civáň et al., 2015), beer yeasts are the products of admixture between different domesticated populations and benefited from historical transfer of fermentation technology.

## Materials and Methods

### Genome sequencing and reference genomes

Genome sequencing was completed for 47 commercial yeast strains, which include 33 ale, 7 lager, 2 whiskey, and 5 baking strains. For reference, sequencing was also completed for 60 strains of diverse origin, including 22 isolates from trees or other non-human-associated sources and 38 isolates from human-associated ferments such as togwa, coffee and cacao (Table S1). For each strain, DNA was extracted and indexed libraries were sequenced on Illumina machines (NextSeq, HiSeq2000 or HiSeq2500). A median of 10.7 million reads per strain was obtained, ranging from 272 thousand to 26 million. The sequencing data is available at NCBI (PRJNA504476).

Genomic data was obtained for 430 strains from publicly available databases. These include 138 additional beer strains from (Gallone et al., 2016; Gonçalves et al., 2016). We also obtained reference genomes for *S. paradoxus*, *S. mikatae* (Scannell et al., 2011), and *S. eubayanus* (SEUB3.0) (Baker et al., 2015). Two large sets of recently published genomes (Duan et al., 2018; Peter et al., 2018) were obtained for comparison with our set of 537 genomes. Genotype calls for SNPs identified in this study were obtained from gvcf files of the 1,011 yeast genomes project (Peter et al., 2018), and genotype calls were generated for 266 strains from China (Duan et al., 2018) using the mapping and genotyping pipeline described below. Because these two later sets of data were only available recently, they were only incorporated into the Figure S1 heatmap.

### Alignment, variant calling and genotyping

Reads were aligned to the *S. cerevisiae* S288c reference genome (R64-1-1_20110203) using BWA-v0.7.12-r1039 (Li, 2013). Lager strains were mapped to a concatenated *S. cerevisiae* and *S. eubayanus* genome and reads mapping to *S. eubayanus* were discarded. For short reads (<70bp) we used BWA-sampe and for the remainder we used BWA-mem. Duplicate reads were marked prior to genotyping. Assembled genomes were also mapped using BWA-mem and flags for secondary alignments were removed to facilitate complete mapping of large contigs. For *S. paradoxus* and *S. mikatae* we obtained higher coverage of the S288c genome by mapping synthetically shredded reads compared to mapping of full contigs and so used the former.

SNPs were called using short read data and then genotyped in those strains with assembled genomes. For SNP calling we used GATK-UnifiedGenotyper-v3.3-0 (Van der Auwera et al., 2013) and applied the hard filters: QD < 5, FS > 60, MQ < 40, MQRankSum < −12.5, and ReadPosRankSum < - 8. The dataset was filtered to remove strains and sites with more than 10% missing data. Among those strains removed were lager strains of the type 1 Saaz group (Dunn and Sherlock, 2008), but we retained *S. paradoxus* and *S. mikatae* for which we obtained calls at 78% and 40% of sites, respectively. Biallelic SNPs with a minor allele frequency of at least 1% and with at least four minor allele genotype calls were selected for analysis, resulting in a total of 273,963 SNPs. The 399 strains retained for analysis are listed in Table S2. Genotype calls for these SNPs were also obtained for the 1,277 strains in the comparative dataset (Duan et al., 2018; Peter et al., 2018).

To estimate our genotyping error rate we compared six pairs of strains that were independently sequenced. Two of the strains, YJF153 and BC217, were haploid derivatives of diploids strains, YPS163 (Doniger et al., 2008) and BC187 (Gerke et al., 2006), respectively, that were also sequenced. The other four pairs were all beer strains independently obtain from Wyeast (Wyeast 1728, 1968, 2565, 2112) and independently sequenced at Washington University in St. Louis and University of Washington in Seattle. Between the pairs of strains we found genotype discordance rates of 9.62×10^-4^ (YJF153/YPS163), 1.31×10^-3^ (BC217/BC187), 3.57×10^-3^ (L.2112/YMD1874), 3.00×10^-3^ (A.2565/YMD1952), 1.81×10^-2^ (A.1968/YMD1981), and 5.74×10^-3^ (A.1728/YMD1866). We retained the six pairs of strains throughout the analysis as a measure of robustness.

### Ploidy and aneuploidy

Ploidy and aneuploidy were assessed by read counts at heterozygous sites and read coverage, respectively. For ploidy analysis, genotypes of 317 strains were from assemblies and so no information on heterozygous sites was available, and 117 strains had few heterozygous sites indicating they were haploid or homozygous diploid. Of the remaining 105 strains, 66 had sufficient coverage at heterozygous sites to make visual designations of ploidy (Forche et al., 2018; Ludlow et al., 2016; Zhu et al., 2016). Visual designations were based on dominant trends consistent with expected percentage of read counts supporting each allele: diploid (50:50), triploid (33:66), tetraploid (25:50:75). Of the 39 strains without sufficient coverage to distinguish triploids from tetraploids, most (33) showed distributions consistent with polyploidy (ploidy > 2), and of these 29 were beer strains (Figure S2). Aneuploidy was assessed by visual inspection of read coverage across the genome. Aneuploidy was only called for clear cases where one or more chromosomes showed a deviation in read coverage compared to all other chromosomes.

### Population structure and admixture

Population structure was inferred by running ADMIXTURE (Alexander et al., 2009) on a set of 20,394 sites with a minimum physical distance of 500 bp. The variants from 138 strains in a recent study of beer strains (Gallone et al., 2016) were removed since the assemblies eliminated heterozygous sites and raw reads for these genomes were not available. Based on 20 independent runs using between 4 and 20 populations for the 399 strains, we chose 13 based on an average change in the log-likelihood greater than 3 standard deviations of the variation in the log-likelihood among independent runs (Figure S3). The beer populations of interest were not affected by this choice; with 12 populations the two Japanese populations merged and with 14 a new population of admixed European wine strains was formed (Figure S3).

Population admixture graphs were inferred using Treemix (Pickrell and Pritchard, 2012). A subset of 199 strains with less than 1% admixture were used to generate a population admixture graph. The population from China was used to root the tree since two strains in the China population, HN6 and SX6 were most closely related to both *S. paradoxus* and *S. mikatae* and blocks of 500 SNPs were used to obtain jacknife standard errors. Five episodes of migration were inferred (P < 4.9e-12), with weights ranging from 0.18-0.49. Migration events were validated using *f_4_* tests of admixture (Table S3). For tests of tree discordance we did not use the clinical and lab populations as reference populations since these showed evidence of admixture. *f_4_* admixture proportions were estimated by the ratio of *f_4_*(Mediterranean, Africa; test, Europe) to *f_4_*(Mediterranean, Africa; Asia, Europe), where each of the 64 beer strains in the Ale 1, Ale 2, lager and beer/baking populations were individually tested.

### Long-read phasing

Three strains were selected for PacBio sequencing and variant phasing. Two of the strains were beer strains, A.2565 and A.T58, and the third, YJF1460, was a hybrid we generated by mating a European/wine strain (BC217) and a Japan/North America 2 oak strain (YJF153). PacBio reads were aligned to the S288c reference genome using Blasr (Chaisson and Tesler, 2012) and heterozygous variants in each genome were phased using HapCUT2 (Edge et al., 2017) and our own heuristic phasing method that accounts for variable ploidy levels across the genome. Average coverage at 56k, 59k and 33k variant sites was 13.1, 18.8 and 13.0 for YJF1460, A.T58, and A.2565, respectively. For our custom phasing method, we used the variant call format files and fragment files from HapCUT2 as input and output a variable number of phased haplotypes. HapCUT2 fragment files were generated with minimum base quality of 10. Reads were merged into haplotypes using a minimum overlap of 4 matching SNPs and a minimum of 80% matching SNPs. Reads were iteratively joined to haplotypes using the best scoring overlap based on score = matches – 5^*^mismatches. Haplotypes were formed by three rounds of merging. In the first round reads were merged into haplotypes without any mismatches. In the second and third rounds haplotypes were merged using the criteria defined above. Error rates were estimated by counting the minimum number of mismatches of reads to the final set of haplotypes. Error rates of 1.84, 2.03 and 1.90% were obtained from comparison of reads to 3337, 2452 and 2607 haplotype alleles for YFJ1460, A.T58 and A.2565, respectively. The average number of haplotypes at phased sites was 2.29, 3.27 and 2.98 for YFJ1460, A.T58 and A.2565, respectively. Sites where three haplotypes were inferred in the YJF1460 control are largely due to overlapping haplotypes that were too short to merge.

After phasing, two sets of SNPs were selected for analysis. The first set consisted of nearly fixed differences between the Europe/wine and Asia/sake populations. After excluding strains with more than 1% admixture, there were 34,022 sites with an allele frequency of 99% in Europe/wine strains (n = 47) and less than 1% frequency in Asia/sake strains (n = 28) or vice-versa. The nearly fixed differences between Europe/wine and Asia/sake strains were used to quantify switching between European and Asian haplotypes. Switching events were measured by counting switches involving 1 or more sites, five or more sites, or sites spanning 4kb or longer (Table S4). The later two measures were used to avoid counting switches caused by sequencing errors or mitotic gene conversion events, which should not affect multiple adjacent sites or regions longer than 4kb (Mancera et al., 2008), respectively. The switching rate for the YJF1460 control was similar to that obtained using HAPCUT2 (Table S4), which minimizes errors when merging reads but assumes a ploidy of two, and SDhaP (Das and Vikalo, 2015) run assuming a ploidy of two for YJF1460 and four for the two ale strains. The second set consisted of alleles abundant in the four beer populations but absent in all others. After excluding strains with more than 1% admixture, there were 32,829 sites with allele frequencies over 25% in either the Ale 1 (n = 13), Ale 2 (n = 12), or Beer/baking strains (n = 2), but less than 1% in all other populations.

We estimated divergence using four-fold degenerate sites in coding sequences. Excluding splice sites and sites with overlapping gene annotations, there were 1,036,317 4-fold degenerate sites surveyed. At these sites, we found 1586, 1040, 899 and 716 alleles at a frequency of 25% or more in the Ale 1, Ale 2, Beer/baking or Lager population, respectively, but not in any other population.

## Acknowledgements

This work was supported by a National Institutes of Health grant (GM080669) to J. Fay, the Rita Allen Foundation, a gift from Karl Handelsman, and a National Science Foundation grant (1516330) to M. Dunham.

---

**Figure S1.**
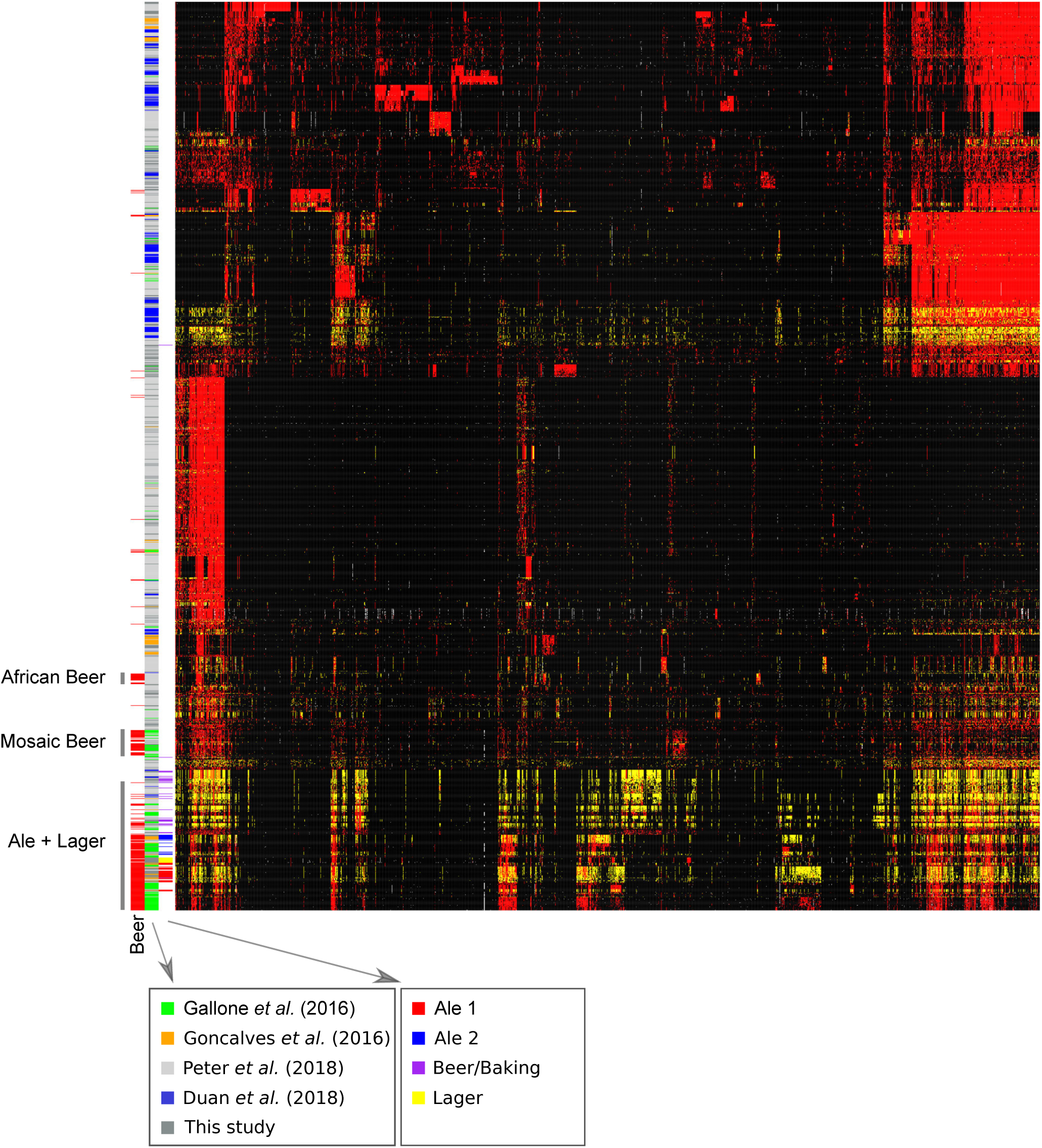
Heatmap of clustered genotypes and strains relating other studies to this one. Strains are color coded by bars, from left to right. Column 1: beer strains (red) with grey labels from Peter et al. (2018) (African Beer, Mosaic Beer) or this study (Ale+Lager); column 2: referenced study; column 3: population assignments from this study. Strains (rows) and SNPs (columns) show genotypes: major allele homozygous (black), heterozygous (yellow), and minor allele homozygous (red).

**Figure S2.**
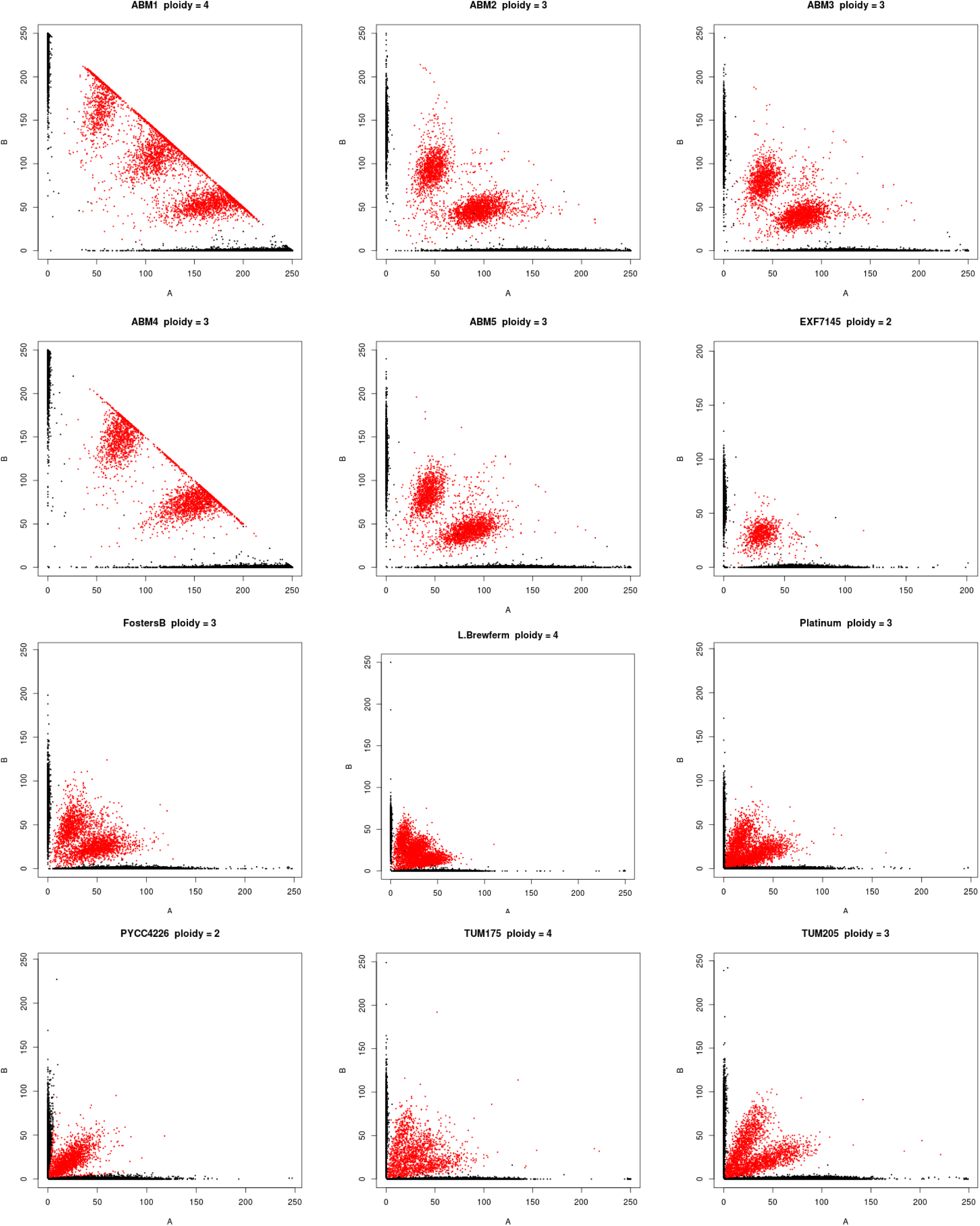
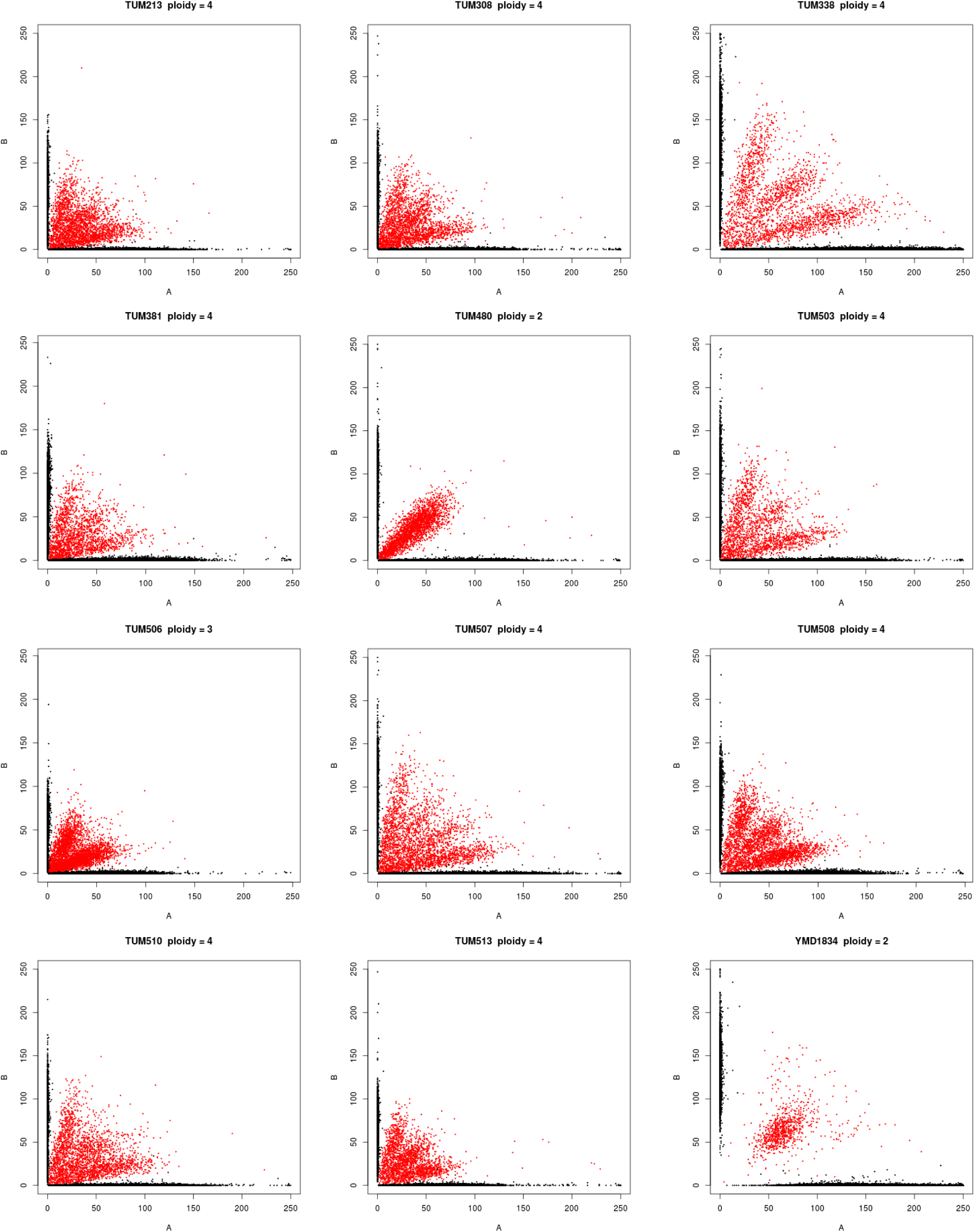
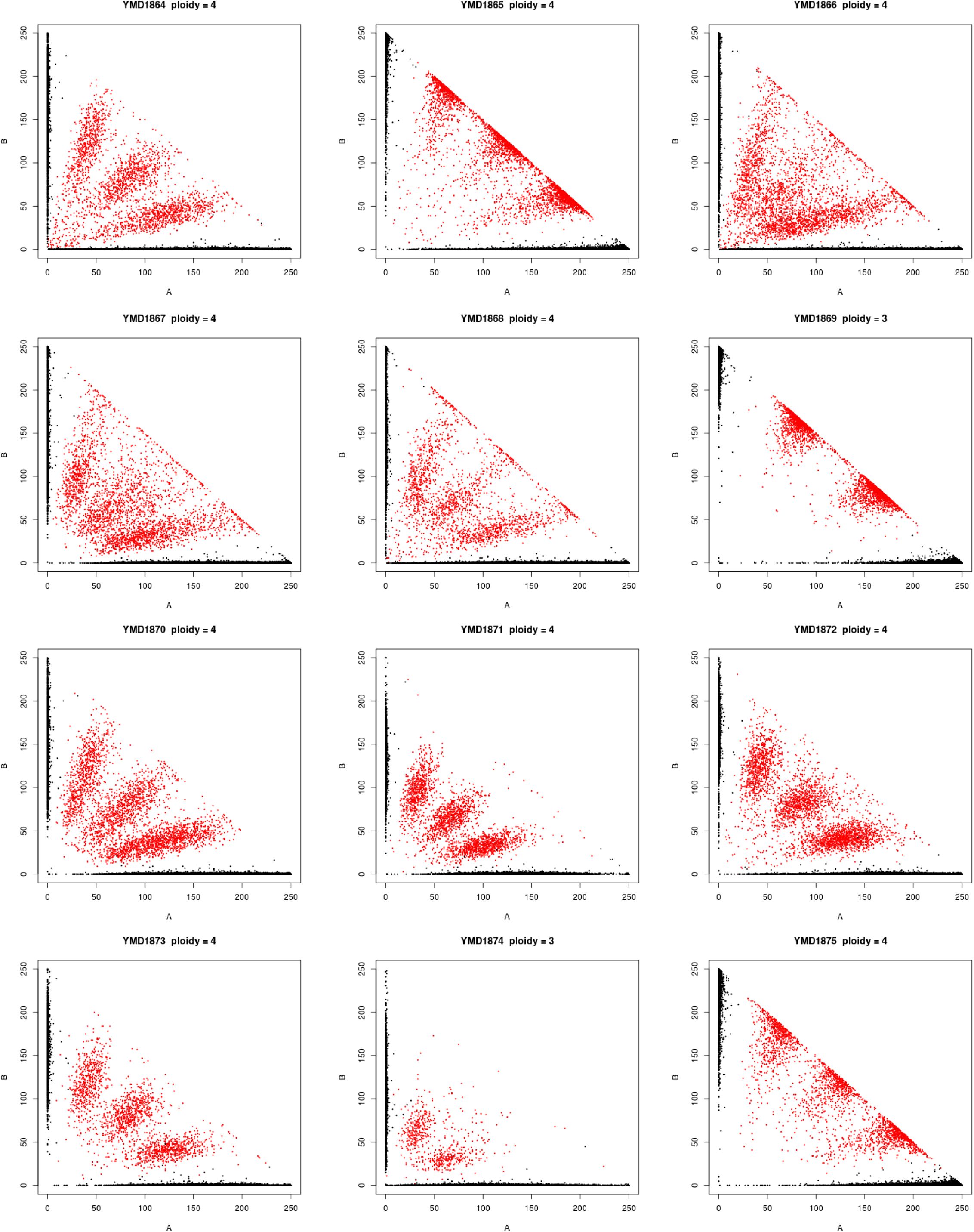
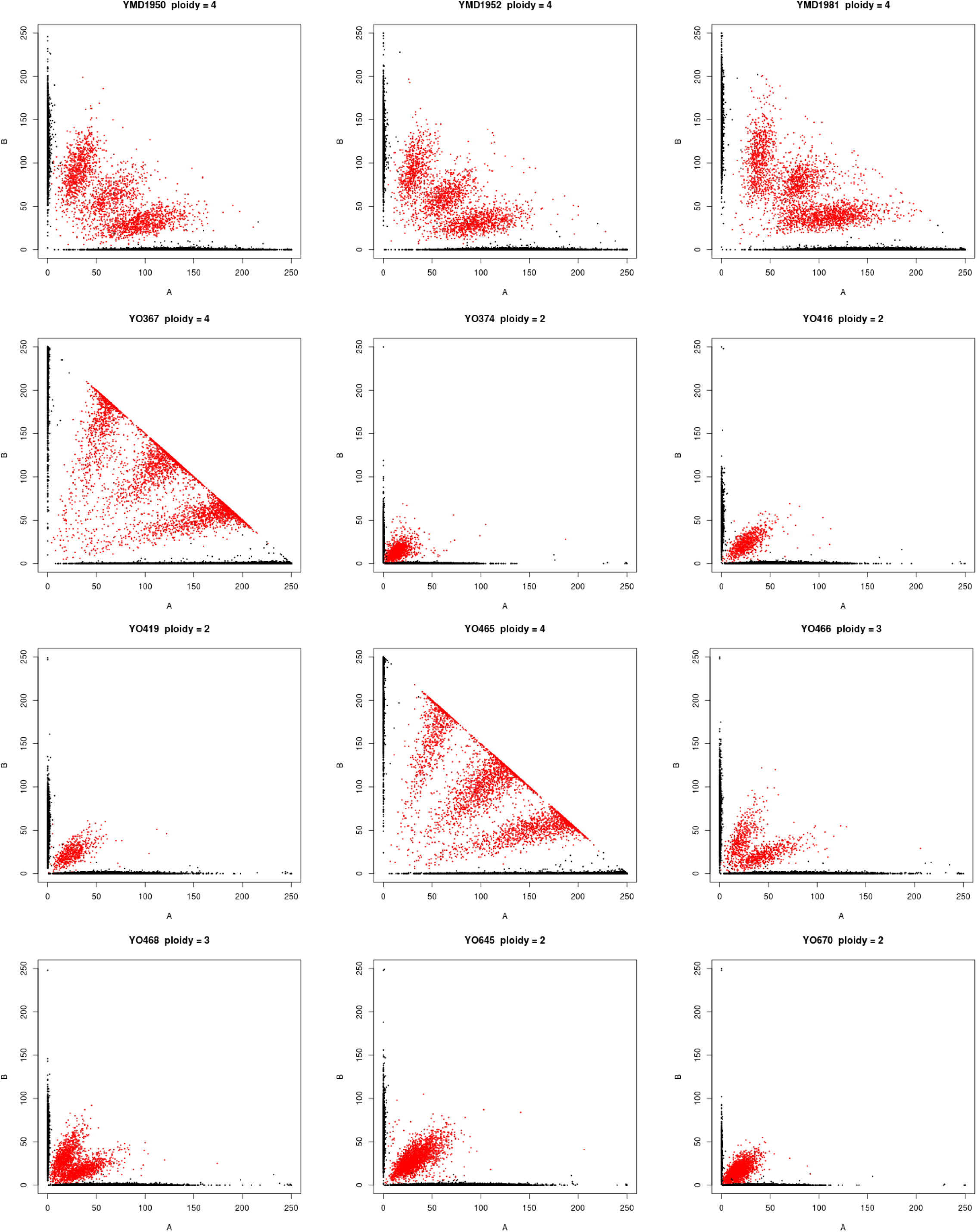
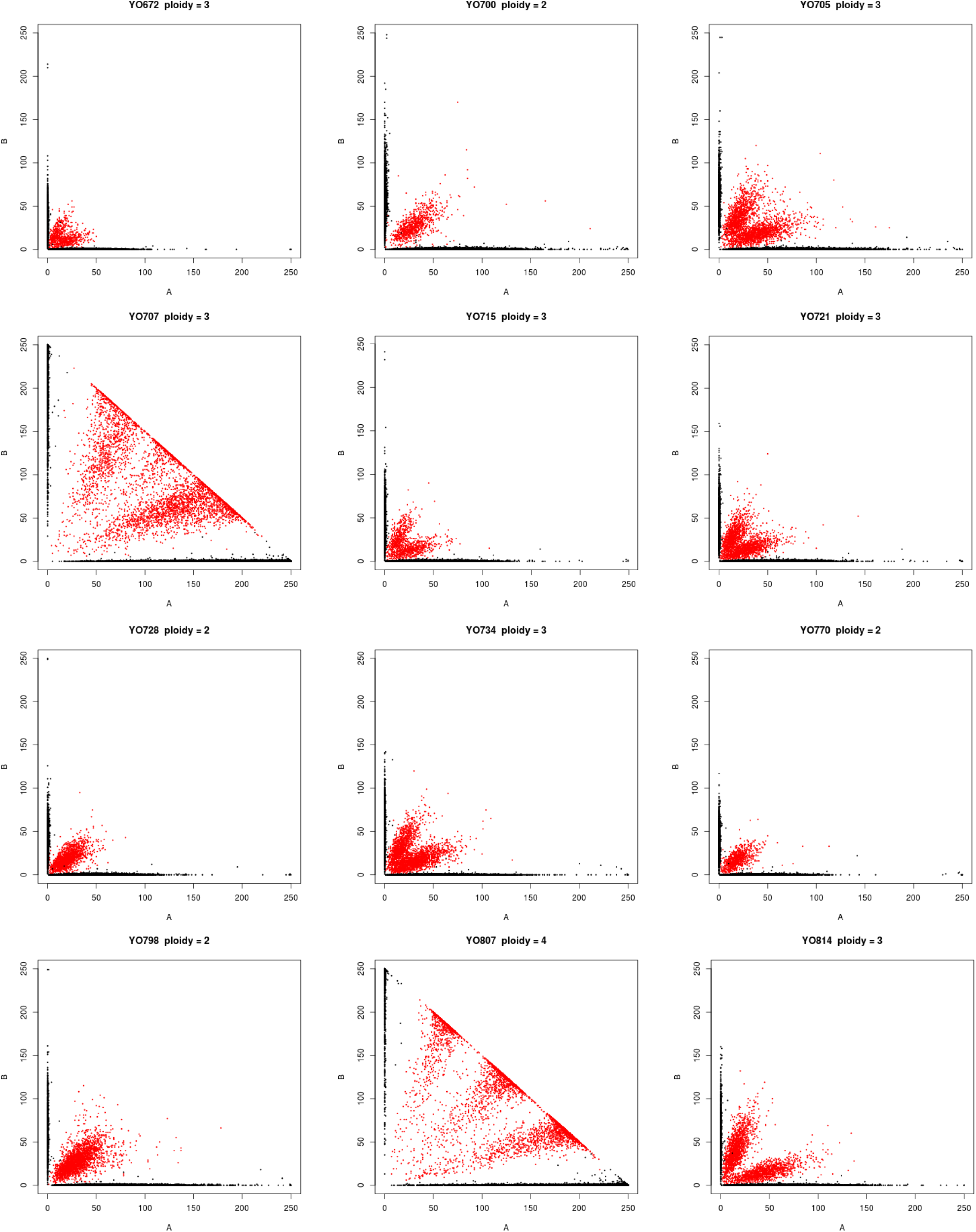
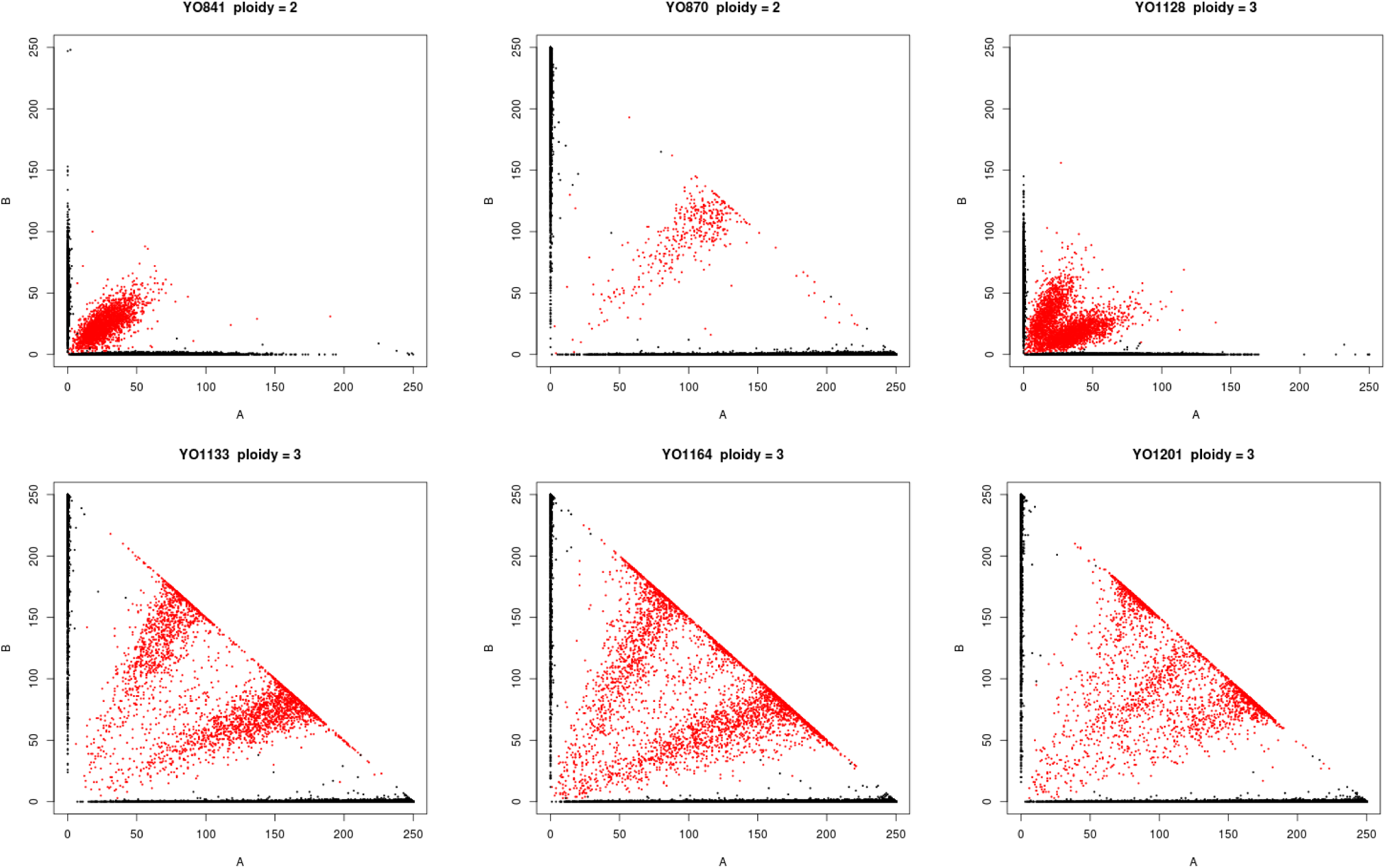
Ploidy inferred from the frequency of genotyped SNPs. Each graph shows the frequency of reads with the reference (A) versus the non-reference (B) allele, with color indicating the genotype call (black = homozygous, red = heterozygous). Diploids, triploids and tetraploids were inferred by heterozygous SNPs being predominantly at frequencies of 50, 33:66 and 25:50:75, respectively.

**Figure S3.**
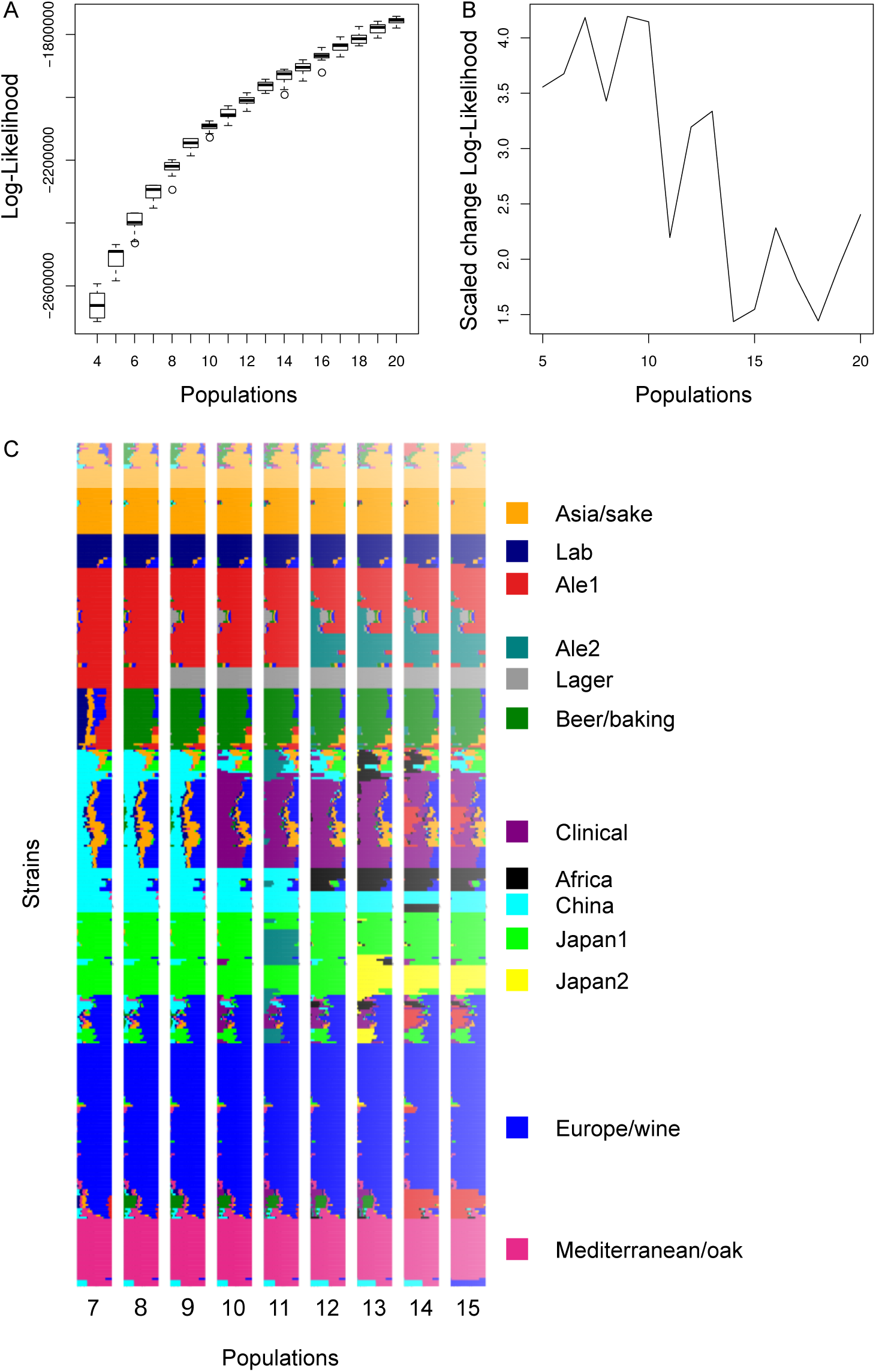
Fit of admixture models as a function of the number of populations. A – Boxplot of the log-likelihood of 20 independent runs as a function of the number of populations. B – The scaled improvement in fit, measured by the change in the log-likelihood with increasing population number divided by the standard deviation in the log-likelihood values from 20 independent runs. C – Population assignments assuming a different number of populations. Each row shows a strain with ancestry to different populations shown by colors and population labels based on similarity to the labels for 13 populations.

**Figure S4.**
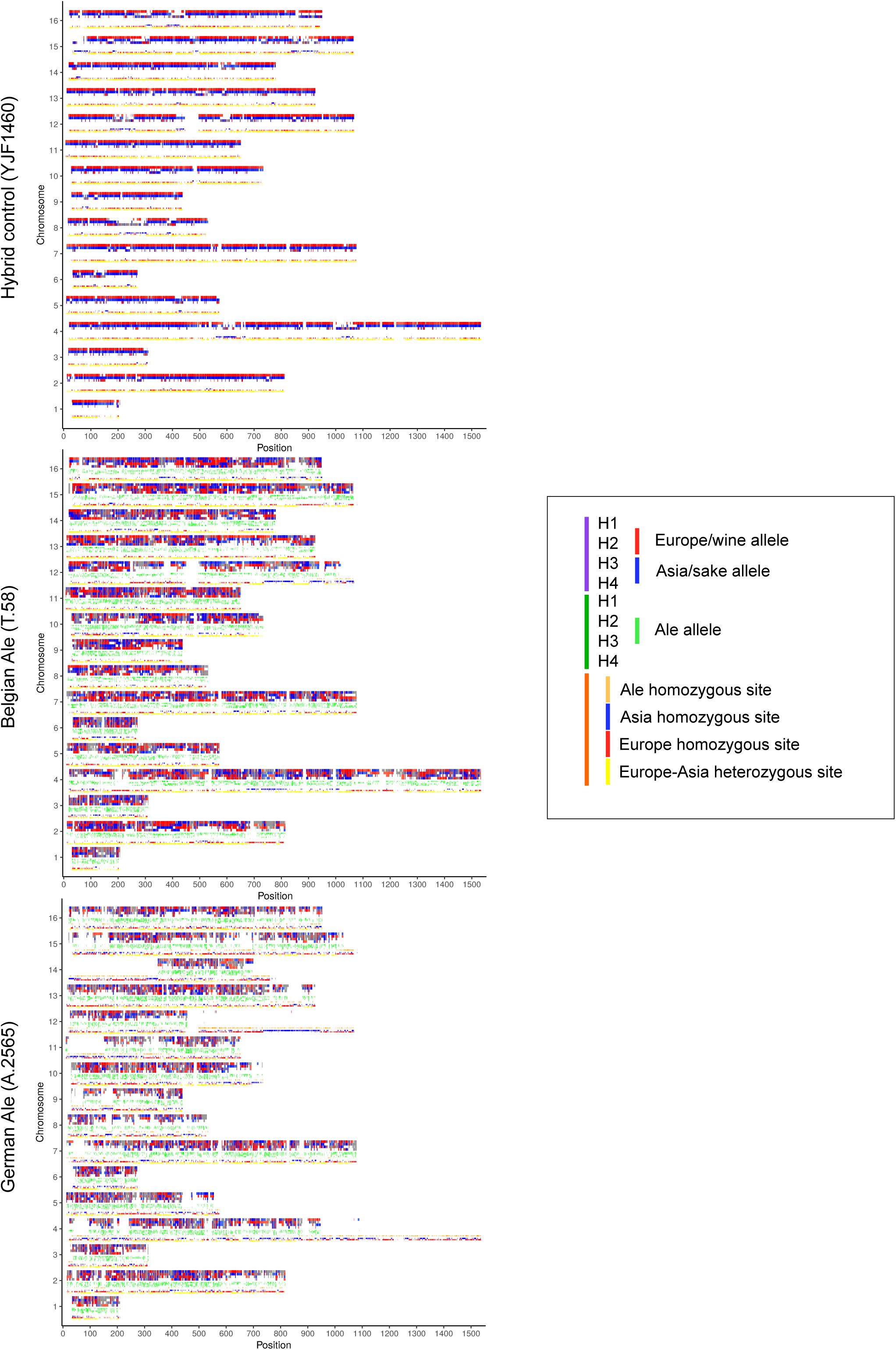
Phased haplotypes show recombination between European and Asian alleles. Panels are the same as in Figure 4 and show all 16 chromosomes for two ale strains (T.58 and A.2565) and the control hybrid (YJF1460). European or Asian alleles are shown in red and blue, respectively, Ale alleles in green. The orange panel shows homozygous Ale, European and Asian alleles as well as sites heterozygous for Europe-Asia alleles.

## References

Alexander, D.H., Novembre, J., and Lange, K. (2009). Fast model-based estimation of ancestry in unrelated individuals. Genome Res. 19, 1655–1664.

Almeida, P., Gonçalves, C., Teixeira, S., Libkind, D., Bontrager, M., Masneuf-Pomarède, I., Albertin, W., Durrens, P., Sherman, D.J., Marullo, P., et al. (2014). A Gondwanan imprint on global diversity and domestication of wine and cider yeast Saccharomyces uvarum. Nat Commun 5, 4044.

Almeida, P., Barbosa, R., Zalar, P., Imanishi, Y., Shimizu, K., Turchetti, B., Legras, J.-L., Serra, M., Dequin, S., Couloux, A., et al. (2015). A population genomics insight into the Mediterranean origins of wine yeast domestication. Mol. Ecol. 24, 5412–5427.

Baker, E., Wang, B., Bellora, N., Peris, D., Hulfachor, A.B., Koshalek, J.A., Adams, M., Libkind, D., and Hittinger, C.T. (2015). The Genome Sequence of Saccharomyces eubayanus and the Domestication of Lager-Brewing Yeasts. Mol. Biol. Evol. 32, 2818–2831.

Bing, J., Han, P.-J., Liu, W.-Q., Wang, Q.-M., and Bai, F.-Y. (2014). Evidence for a Far East Asian origin of lager beer yeast. Curr. Biol. 24, R380-381.

Campbell, D., Doctor, J.S., Feuersanger, J.H., and Doolittle, M.M. (1981). Differential mitotic stability of yeast disomes derived from triploid meiosis. Genetics 98, 239–255.

Chaisson, M.J., and Tesler, G. (2012). Mapping single molecule sequencing reads using basic local alignment with successive refinement (BLASR): application and theory. BMC Bioinformatics 13, 238.

Choi, J.Y., Platts, A.E., Fuller, D.Q., Hsing (邢禹依), Y.-I., Wing, R.A., and Purugganan, M.D. (2017). The Rice Paradox: Multiple Origins but Single Domestication in Asian Rice. Mol Biol Evol 34, 969–979.

Civáň, P., Craig, H., Cox, C.J., and Brown, T.A. (2015). Three geographically separate domestications of Asian rice. Nat Plants 1, 15164.

Cromie, G.A., Hyma, K.E., Ludlow, C.L., Garmendia-Torres, C., Gilbert, T.L., May, P., Huang, A.A., Dudley, A.M., and Fay, J.C. (2013). Genomic sequence diversity and population structure of Saccharomyces cerevisiae assessed by RAD-seq. G3 (Bethesda) 3, 2163–2171.

Das, S., and Vikalo, H. (2015). SDhaP: haplotype assembly for diploids and polyploids via semi-definite programming. BMC Genomics 16, 260.

Dashko, S., Zhou, N., Compagno, C., and Piškur, J. (2014). Why, when, and how did yeast evolve alcoholic fermentation? FEMS Yeast Res. 14, 826–832.

Dashko, S., Liu, P., Volk, H., Butinar, L., Piškur, J., and Fay, J.C. (2016). Changes in the Relative Abundance of Two Saccharomyces Species from Oak Forests to Wine Fermentations. Front Microbiol 7, 215.

Doniger, S.W., Kim, H.S., Swain, D., Corcuera, D., Williams, M., Yang, S.-P., and Fay, J.C. (2008). A catalog of neutral and deleterious polymorphism in yeast. PLoS Genet 4, e1000183.

Duan, S.-F., Han, P.-J., Wang, Q.-M., Liu, W.-Q., Shi, J.-Y., Li, K., Zhang, X.-L., and Bai, F.-Y. (2018). The origin and adaptive evolution of domesticated populations of yeast from Far East Asia. Nat Commun 9, 2690.

Dunn, B., and Sherlock, G. (2008). Reconstruction of the genome origins and evolution of the hybrid lager yeast Saccharomyces pastorianus. Genome Res. 18, 1610–1623.

Edge, P., Bafna, V., and Bansal, V. (2017). HapCUT2: robust and accurate haplotype assembly for diverse sequencing technologies. Genome Res. 27, 801–812.

Fay, J.C., and Benavides, J.A. (2005). Evidence for domesticated and wild populations of Saccharomyces cerevisiae. PLoS Genetics 1, 66–71.

Forche, A., Cromie, G., Gerstein, A.C., Solis, N.V., Pisithkul, T., Srifa, W., Jeffery, E., Abbey, D., Filler, S.G., Dudley, A.M., et al. (2018). Rapid Phenotypic and Genotypic Diversification After Exposure to the Oral Host Niche in Candida albicans. Genetics 209, 725–741.

Gallone, B., Steensels, J., Prahl, T., Soriaga, L., Saels, V., Herrera-Malaver, B., Merlevede, A., Roncoroni, M., Voordeckers, K., Miraglia, L., et al. (2016). Domestication and Divergence of Saccharomyces cerevisiae Beer Yeasts. Cell 166, 1397-1410.e16.

Gerke, J., Lorenz, K., and Cohen, B. (2009). Genetic interactions between transcription factors cause natural variation in yeast. Science 323, 498–501.

Gerke, J.P., Chen, C.T.L., and Cohen, B.A. (2006). Natural isolates of Saccharomyces cerevisiae display complex genetic variation in sporulation efficiency. Genetics 174, 985–997.

Gonçalves, M., Pontes, A., Almeida, P., Barbosa, R., Serra, M., Libkind, D., Hutzler, M., Gonçalves, P., and Sampaio, J.P. (2016). Distinct Domestication Trajectories in Top-Fermenting Beer Yeasts and Wine Yeasts. Curr. Biol. 26, 2750–2761.

González, S.S., Barrio, E., Gafner, J., and Querol, A. (2006). Natural hybrids from Saccharomyces cerevisiae, Saccharomyces bayanus and Saccharomyces kudriavzevii in wine fermentations. FEMS Yeast Res. 6, 1221–1234.

Green, R.E., Krause, J., Briggs, A.W., Maricic, T., Stenzel, U., Kircher, M., Patterson, N., Li, H., Zhai, W., Fritz, M.H.-Y., et al. (2010). A draft sequence of the Neandertal genome. Science 328, 710–722.

Hagman, A., Säll, T., Compagno, C., and Piskur, J. (2013). Yeast “make-accumulate-consume” life strategy evolved as a multi-step process that predates the whole genome duplication. PLoS One 8, e68734.

Hyma, K.E., and Fay, J.C. (2013). Mixing of vineyard and oak-tree ecotypes of Saccharomyces cerevisiae in North American vineyards. Mol. Ecol. 22, 2917–2930.

Lang, G.I., and Murray, A.W. (2008). Estimating the Per-Base-Pair Mutation Rate in the Yeast Saccharomyces cerevisiae. Genetics 178, 67–82.

Lee, P.S., Greenwell, P.W., Dominska, M., Gawel, M., Hamilton, M., and Petes, T.D. (2009). A fine-structure map of spontaneous mitotic crossovers in the yeast Saccharomyces cerevisiae. PLoS Genet. 5, e1000410.

Legras, J.-L., Merdinoglu, D., Cornuet, J.-M., and Karst, F. (2007). Bread, beer and wine: Saccharomyces cerevisiae diversity reflects human history. Mol. Ecol. 16, 2091–2102.

Legras, J.-L., Erny, C., and Charpentier, C. (2014). Population structure and comparative genome hybridization of European flor yeast reveal a unique group of Saccharomyces cerevisiae strains with few gene duplications in their genome. PLoS ONE 9, e108089.

Legras, J.-L., Galeote, V., Bigey, F., Camarasa, C., Marsit, S., Nidelet, T., Sanchez, I., Couloux, A., Guy, J., Franco-Duarte, R., et al. (2018). Adaptation of S. cerevisiae to Fermented Food Environments Reveals Remarkable Genome Plasticity and the Footprints of Domestication. Mol. Biol. Evol. 35, 1712–1727.

Li, H. (2013). Aligning sequence reads, clone sequences and assembly contigs with BWA-MEM. ArXiv:1303.3997 [q-Bio].

Libkind, D., Hittinger, C.T., Valério, E., Gonçalves, C., Dover, J., Johnston, M., Gonçalves, P., and Sampaio, J.P. (2011). Microbe domestication and the identification of the wild genetic stock of lager-brewing yeast. Proc. Natl. Acad. Sci. U.S.A. 108, 14539–14544.

Liti, G., Carter, D.M., Moses, A.M., Warringer, J., Parts, L., James, S.A., Davey, R.P., Roberts, I.N., Burt, A., Koufopanou, V., et al. (2009). Population genomics of domestic and wild yeasts. Nature 458, 337–341.

Loidl, J. (1995). Meiotic chromosome pairing in triploid and tetraploid Saccharomyces cerevisiae. Genetics 139, 1511–1520.

Ludlow, C.L., Cromie, G.A., Garmendia-Torres, C., Sirr, A., Hays, M., Field, C., Jeffery, E.W., Fay, J.C., and Dudley, A.M. (2016). Independent Origins of Yeast Associated with Coffee and Cacao Fermentation. Curr. Biol. 26, 965–971.

Mallet, J. (2007). Hybrid speciation. Nature 446, 279–283.

Mancera, E., Bourgon, R., Brozzi, A., Huber, W., and Steinmetz, L.M. (2008). High-resolution mapping of meiotic crossovers and non-crossovers in yeast. Nature 454, 479–485.

McGovern, P.E. (2013). Ancient Wine: The Search for the Origins of Viniculture (Princeton University Press).

Myles, S., Boyko, A.R., Owens, C.L., Brown, P.J., Grassi, F., Aradhya, M.K., Prins, B., Reynolds, A., Chia, J.-M., Ware, D., et al. (2011). Genetic structure and domestication history of the grape. PNAS 108, 3530–3535.

Nakazawa, N., Ashikari, T., Goto, N., Amachi, T., Nakajima, R., Harashima, S., and Oshima, Y. (1992). Partial restoration of sporulation defect in sake yeasts, kyokai no. 7 and no. 9, by increased dosage of the IME1 gene. Journal of Fermentation and Bioengineering 73, 265–270.

Peris, D., Langdon, Q.K., Moriarty, R.V., Sylvester, K., Bontrager, M., Charron, G., Leducq, J.-B., Landry, C.R., Libkind, D., and Hittinger, C.T. (2016). Complex Ancestries of Lager-Brewing Hybrids Were Shaped by Standing Variation in the Wild Yeast Saccharomyces eubayanus. PLoS Genet. 12, e1006155.

Peter, B.M. (2016). Admixture, Population Structure, and F-Statistics. Genetics 202, 1485–1501.

Peter, J., De Chiara, M., Friedrich, A., Yue, J.-X., Pflieger, D., Bergström, A., Sigwalt, A., Barre, B., Freel, K., Llored, A., et al. (2018). Genome evolution across 1,011 Saccharomyces cerevisiae isolates. Nature 556, 339–344.

Pickrell, J.K., and Pritchard, J.K. (2012). Inference of population splits and mixtures from genome-wide allele frequency data. PLoS Genet. 8, e1002967.

Poets, A.M., Fang, Z., Clegg, M.T., and Morrell, P.L. (2015). Barley landraces are characterized by geographically heterogeneous genomic origins. Genome Biology 16, 173.

Reich, D., Thangaraj, K., Patterson, N., Price, A.L., and Singh, L. (2009). Reconstructing Indian population history. Nature 461, 489–494.

Salman-Minkov, A., Sabath, N., and Mayrose, I. (2016). Whole-genome duplication as a key factor in crop domestication. Nat Plants 2, 16115.

Scannell, D.R., Zill, O.A., Rokas, A., Payen, C., Dunham, M.J., Eisen, M.B., Rine, J., Johnston, M., and Hittinger, C.T. (2011). The Awesome Power of Yeast Evolutionary Genetics: New Genome Sequences and Strain Resources for the Saccharomyces sensu stricto Genus. G3 (Bethesda) 1, 11–25.

Schacherer, J., Shapiro, J.A., Ruderfer, D.M., and Kruglyak, L. (2009). Comprehensive polymorphism survey elucidates population structure of Saccharomyces cerevisiae. Nature 458, 342–345.

Sipiczki, M. (2008). Interspecies hybridization and recombination in Saccharomyces wine yeasts. FEMS Yeast Res. 8, 996–1007.

Sniegowski, P.D., Dombrowski, P.G., and Fingerman, E. (2002). Saccharomyces cerevisiae and Saccharomyces paradoxus coexist in a natural woodland site in North America and display different levels of reproductive isolation from European conspecifics. FEM Yeast Res 1, 299–306.

Strope, P.K., Skelly, D.A., Kozmin, S.G., Mahadevan, G., Stone, E.A., Magwene, P.M., Dietrich, F.S., and McCusker, J.H. (2015). The 100-genomes strains, an S. cerevisiae resource that illuminates its natural phenotypic and genotypic variation and emergence as an opportunistic pathogen. Genome Res. 25, 762–774.

Tilakaratna, V., and Bensasson, D. (2017). Habitat Predicts Levels of Genetic Admixture in Saccharomyces cerevisiae. G3 (Bethesda) 7, 2919–2929.

Van de Peer, Y., Mizrachi, E., and Marchal, K. (2017). The evolutionary significance of polyploidy. Nat. Rev. Genet. 18, 411–424.

Van der Auwera, G.A., Carneiro, M.O., Hartl, C., Poplin, R., Del Angel, G., Levy-Moonshine, A., Jordan, T., Shakir, K., Roazen, D., Thibault, J., et al. (2013). From FastQ data to high confidence variant calls: the Genome Analysis Toolkit best practices pipeline. Curr Protoc Bioinformatics 43, 11.10.1-33.

Wang, Q.-M., Liu, W.-Q., Liti, G., Wang, S.-A., and Bai, F.-Y. (2012). Surprisingly diverged populations of Saccharomyces cerevisiae in natural environments remote from human activity. Mol. Ecol. 21, 5404–5417.

Williams, K.M., Liu, P., and Fay, J.C. (2015). Evolution of ecological dominance of yeast species in high-sugar environments. Evolution 69, 2079–2093.

Yim, E., O’Connell, K.E., St Charles, J., and Petes, T.D. (2014). High-resolution mapping of two types of spontaneous mitotic gene conversion events in Saccharomyces cerevisiae. Genetics 198, 181–192.

Zhu, Y.O., Sherlock, G., and Petrov, D.A. (2016). Whole Genome Analysis of 132 Clinical Saccharomyces cerevisiae Strains Reveals Extensive Ploidy Variation. G3 (Bethesda) 6, 2421–2434.

